# Cooperation increases robustness to ecological disturbance in microbial cross-feeding networks

**DOI:** 10.1101/2020.05.15.098103

**Authors:** Leonardo Oña, Christian Kost

## Abstract

Microorganisms mainly exist within complex networks of ecological interactions. Given that the growth and survival of community members frequently depend on an obligate exchange of essential metabolites, it is generally unclear how such communities can persist despite the destabilizing force of ecological disturbance. Here we address this issue using a population dynamics model. In contrast to previous work that suggests the potential for obligate interaction networks to evolve is limited, we find the opposite pattern: natural selection in the form of ecological disturbance favors both specific network topologies and cooperative cross-feeding among community members. These results establish environmental selection as a key driver shaping the architecture of microbial interaction networks.

## 1 Significance

Microbes live in diverse communities that significantly impact the ecology and evolution of other organisms. These communities represent complex ecological networks, within which the constituent strains engage in an obligate exchange of essential metabolites. However, it remains generally unclear how these interactions can persist in the face of strong ecological perturbation. Here, we address this question using a population dynamics model. We show that both the topology of the interaction network and the degree to which microbes engage in a cooperative exchange of metabolites shape the systems robustness to ecological disturbance. Thus, our study identifies key principles underlying metabolic trade in natural microbial communities. These results can help to design synthetic microbial consortia for medical and biotechnological applications.

## 2 Introduction

Microbial communities play key roles in many ecosystems^1,2^ and contribute significantly to the maintenance of plant and animal health^3,4,5^. In most cases, these assemblages consist of a large number of metabolically diverse genotypes that engage in a complex network of both antagonistic and synergistic ecological interactions^6,7^. While it is clear that the interplay between these different interactions determines the structure, function, and evolution of a given microbial community, the general principles guiding this process remain poorly understood. However, a detailed knowledge of how properties of individual strains combine to give rise to emergent phenotypes at the community-level is not only central to our understanding of microbial ecology, but also to the design of synthetic microbial communities for medical or biotechnological applications^8,6^.

One specific type of ecological interaction that appears to be particularly important in microbial communities is the exchange of essential metabolites among community members^9,10,11^. A growing body of literature suggests that a large proportion of all bacteria known lacks biosynthetic pathways to autonomously produce essential building block metabolites such as amino acids, vitamins, and even nucleotides^12,13,14^. Thus, growth and survival of these so-called *auxotrophic* microorganisms depends on the presence of other individuals that provide sufficient amounts of the required metabolites. The evolutionary process that likely drives the emergence of such metabolic dependencies has been termed *Black Queen* dynamics^15,16^. The basic idea is that as microbes grow, they commonly release significant amounts of metabolites in the extracellular environment. These compounds represent a valuable resource (i.e., a so-called *public good*) that can be used by newly arising auxotrophic genotypes that lack the ability to autonomously produce the corresponding metabolites. In this way, an obligate metabolic interaction is generated that ties the fate of the auxotrophic recipient to the presence of other cells that can provide it with the required metabolite. Interestingly, it has been shown that auxotrophic mutants gain a significant fitness advantage from using external metabolite sources, because they save the energy to produce the compound on their own^14,16^. This type metabolic cross-feeding interaction, which initially relies on an exchange of metabolic by-products, can be further strengthened when auxotrophic genotypes start to increase the production level of the traded chemical^17,18,19^. Such an increased investment that is costly to the producing cell, can be favored by natural selection, when the cooperative cell is rewarded for its initiative by receiving fitness benefits in return. This can be the case when two types interact that reciprocally exchange essential metabolites^17,19^ or if the interaction is staged in a spatially structured environment with low rates of population intermixing^20,21^. In the long-run, this evolutionary process is expected to give rise to multipartite microbial networks of different sizes and topologies, within which metabolites are reciprocally exchanged^10^.

In the beginning, such interaction networks are likely created by chance: resident auxotrophic mutants and prototrophic genotypes that share the same environment start engaging in metabolic cross-feeding interactions. Depending on the amounts of metabolites the interacting cells produce and consume, the resulting interaction collapses immediately or remains stable for extended periods. However, what determines the stability of these highly-interwoven interaction networks? Given that the survival of auxotrophic cells critically depends on the presence of other individuals that can provide the required metabolite, loss of these donor cells may lead to a catastrophic collapse of the entire microbial community. Indeed, a previous theoretical study on the ecological stability of microbial community networks concluded that cooperating networks of microbes are often unstable^22^. In this study, stability was modeled as resilience: the capacity to return to the equilibrium after a transient change in population size. However, changes in the environment often cause modifications of population growth rates, instead of merely a transient decrease in the number of individuals. Moreover, given that the public goods that are exchanged between microbial strains are often key for determining community stability, their dynamics should be taken into account as well^23^. Furthermore, even though public goods can negatively affect yield and stability of microbial communities when they are produced in sub-optimal concentrations, they can also positively affect these parameters when they are efficiently produced and involved in an effect called ‘division of labor’. Here, two or more strains can save energy by distributing the production of certain metabolites among the participating individuals and subsequently exchanging the produced compounds^18,24^. However, this division of labor effect was not considered in previous models^22^. Taken together, environmental variables can directly and indirectly affect population growth rates by modifying the production and availability of public goods. In this way, environmental changes can impinge upon the dynamics and stability of microbial communities.

Changes in environmental variables are more explicitly incorporated in models as ecological disturbance by modifying one or several parameters that affect the growth rate of the focal population. Disturbance, in the form of periodically-occurring or constant perturbations of the environment, can cause mass-mortality and has been shown to strongly affect the composition, structure, and function of microbial communities from diverse habitats such as soil^25^, lakes^26^, and the human gut^27,28^. Models incorporating disturbance in microbial communities revealed that the response to such perturbations can exhibit complex dynamics^29^. Following disturbance, the community can, for example, undergo critical transitions to alternative stable states^30,29^ or approach a catastrophic collapse of the entire community^31,32^. The architecture of the network that is determined by the ecological interactions among individual types can strongly affect the robustness of biological communities to such events of ecological disturbance^33^. This is the case, for example, for pollination, seed dispersal^34^, and trophic networks^35^. However, very little is known on how ecological networks of different sizes and topologies within microbial communities respond to disturbance^13^.

Here we fill this gap by using a population dynamics model to analyze networks of auxotrophic microorganisms that exchange metabolites as extracellular public goods. In particular, we aim at identifying how the robustness of these networks to ecological disturbance is affected by: (i) the number of auxotrophy-causing mutations, (ii) the topology of the interaction network, and (iii) the presence of cooperative cross-feeders within the network. Our analysis revealed that, all else being equal, communities with more auxotrophy-causing mutations were less robust to disturbance. Second, the network topology of metabolite production strongly affected the system’s stability. Finally, mutations that increased amino acid production levels of auxotrophic microbes within interaction networks increased the robustness of these communities to ecological disturbance.

## 3 Methods

The dynamics of *n* auxotrophic microbes (*B*_*i*_) and *m* metabolites (*M*_*k*_) is described by the following system of ordinary differential equations:

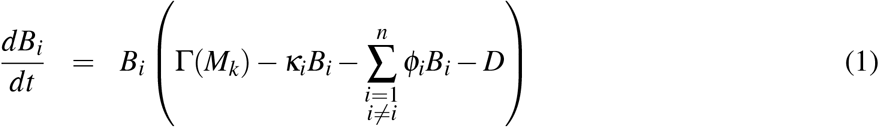

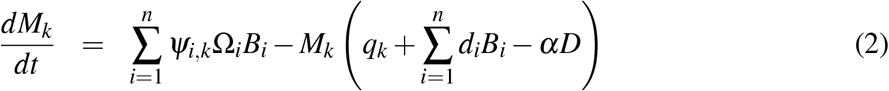

The function Γ(*M*_*k*_) takes different forms depending on the model assumptions (see Supplementary Material for details and extensions). In the model used throughout the main text, this function is given by 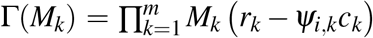, where the per capita growth of microbes is the result of the utilization of all metabolites with *r*_*k*_ denoting the per capita growth rate and *c*_*k*_ is the cost associated with the production for the *k*^*th*^ metabolite. The obligate nature of the interaction between microbes and metabolites is represented by the term 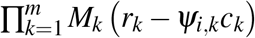. This product ensures that, when at least one of the metabolites *M*_*k*_ is zero, all microbes are going extinct. The terms *κ*_*i*_ and *ϕ*_*i*_ are the rate of intra- and inter-specific competition. Microbes produce metabolites at a rate Ω_*i*_ and the intake of metabolites occurs at a rate *d*_*i*_. Metabolites are also assumed to be lost by degradation or diffusion into the environment at a rate *q*_*k*_. The parameter *ψ*_*i,k*_ represent the presence/absence of a mutation causing auxotrophy, and therefore can take the values 0 or 1, defining the network of auxotrophs. In the main model, we assumed that metabolites themselves are not affected by disturbance (*α* = 0). However, relaxing this assumption did not change the results (see the Supplementary Material).

### 3.1 Disturbance in random networks of auxotrophs

Simulations are based on different parametrization of the model given by equations (1, 2), describing the microbial dynamics and the metabolites produced. We created microbial systems assuming that all parameters affecting the dynamics are the same among microbes, and the same among metabolites (in the main text) and vary only the position of mutation-causing auxotrophies in the network. This allowed us to study the stability of the resulting network topology of metabolite production in isolation, without the confounding effect that would result if parameters were different among microbes and metabolites. To ensure that our results still hold when this assumption is relaxed, we assigned random parameters for microbes and metabolites in another set of simulations (see Supplementary Material).

In all cases, network topologies are given by a bipartite graph describing metabolite production by microbes (Fig.1a). Here, a network is formalized by a matrix in which entries contain the production of the *k*^*th*^ metabolite by the *i*^*th*^ microbe. A microbial system of prototrophs producing metabolites is given by a matrix of size *n x m*, where all *ψ*_*i,k*_ = 1. In such a microbial system, a mutation causing auxotrophy is symbolized by a particular entry in the matrix where *ψ*_*i,k*_ = 0. For a given fixed number of auxotrophy-causing mutations, different matrices are randomly generated, characterized by a fixed number of zeros in the entries, but in different positions in the matrix. This diversity of patterns in the randomly generated matrices, under the formalization of the network theory, was characterized by different topologies. Not all topologies resulting from the randomization process generate microbial systems with a stable equilibrium, where all microbes and all metabolites are non-zero. Thus, we only incorporated cases, where the resulting system of equations has a stable equilibrium with non-zero microbes and metabolites, and such equilibrium is stable (i.e., all eigenvalues were negative). For this, we first assumed that *D* = 0, then obtained the equilibrium where all variables were non-zero, and finally used those values as the initial condition for a new set of ordinary differential equations, where the disturbance term was added to the equation for the microbes (*D* > 0). This type of disturbance has been called ‘press disturbance’^36^ (see Supplementary Material for details). Each microbial system generated through the randomization process was exposed to increased levels of disturbance (i.e., increasing the numerical value of parameter *D*) (Fig.1b) until it went extinct. The disturbance value *D*, where the first extinction took place, was defined as the robustness for that microbial system.

**Fig. 1.**
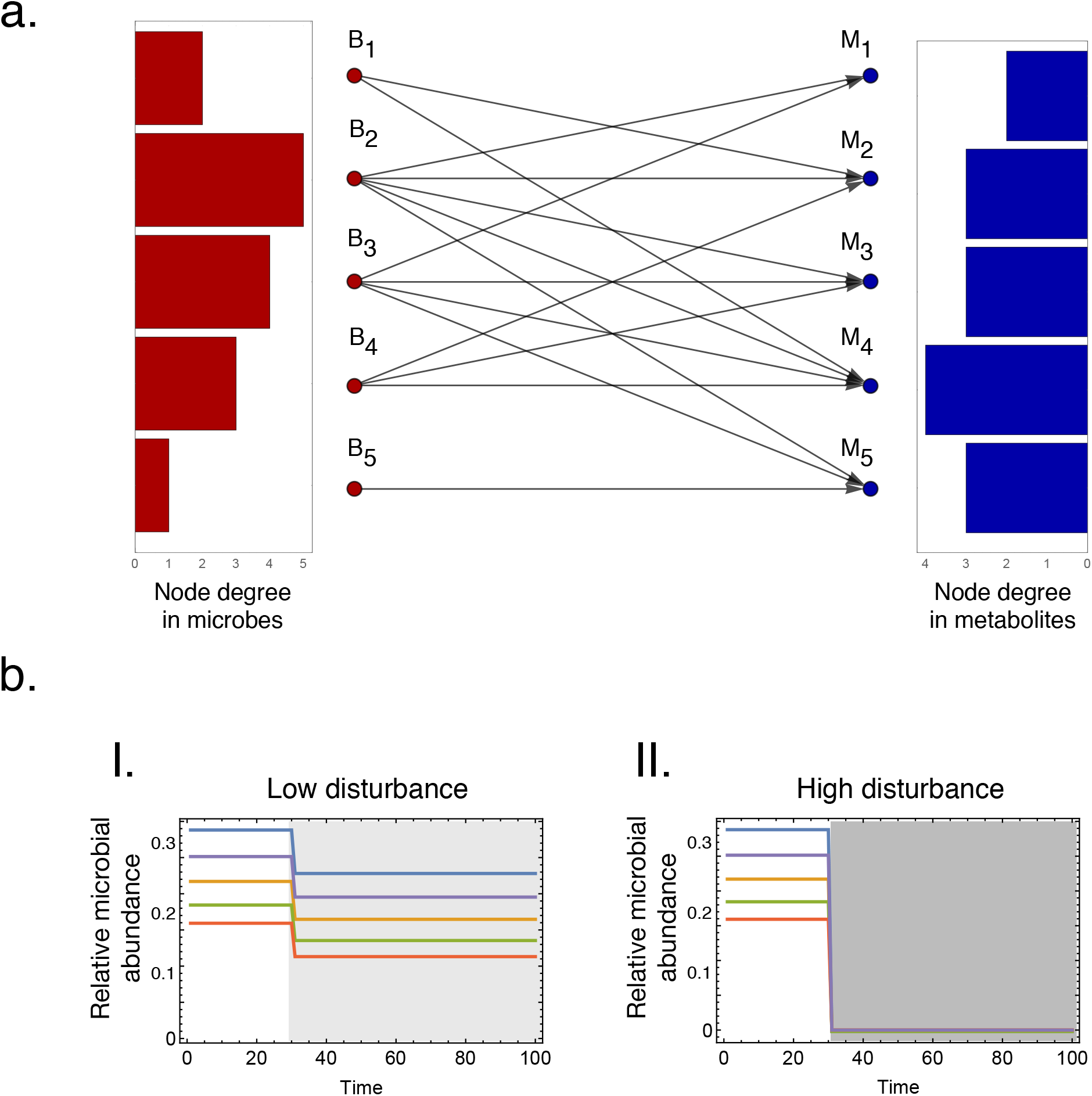
Networks of auxotrophic microorganisms and the effect of environmental disturbance. **(a)** Exemplary network depicting microorganisms *B*_1_ *B*_5_ that produce metabolites *M*_1_ *M*_5_. We define the *auxotrophy degree in microbes*, as the number of metabolites a microbe is unable to produce (i.e., the inverse of the node degree). In the example, *B*_1_ has an auxotrophy degree of 3, while *B*_5_ has an auxotrophy degree of 4. In a similar way, we define the *auxotrophy degree in metabolites*, as the number of microbes that are unable to produce a certain metabolite. In the example, *M*_1_ has a degree of 3, while *M*_4_ has a degree of 1. These quantities, which are shown as a bar plot, are important to characterize the topology of the network calculating the *normalized entropy* (i.e., using the distribution of the auxotrophy degree in microbes) or the *assortativity* of the network (i.e., using the distribution of the auxotrophy degree in both microbes and metabolites) (see main text for an explanation of these terms). **(b)** Effect of ecological disturbance on the dynamics of the ecological community shown in Fig.1a. The network was disturbed with (I.) a low (*D* = 70) or (II.) a high (*D* = 136) intensity (i.e., shadowed areas).

### 3.2 Normalized entropy and assortativity

Each randomly generated microbial system is represented by an interaction network between microbes and metabolites with a certain topology, which can be depicted as a bipartite network (Fig.1a). Analyzing these networks, we aimed at finding topological properties that correlate with robust responses to disturbance. Two measures, which describe the degree of homogeneity with which metabolite production is distributed among microbes, were strongly correlated with network robustness. The first one defines how evenly auxotrophy-causing mutation are distributed in the microbial community. The corresponding value is simply given by the entropy of the distribution of mutations causing auxotrophy, relative to the maximum entropy possible for that particular number of mutations, and it is called the ‘normalized entropy’ (*node degree in microbes*, given by the red bar plot in Fig.1a). On the other hand, metabolites can be produced by a different number of microbes (*node degree of metabolites*, given by the blue bar plot in Fig.1a. This measure describes how evenly the production of each metabolite is distributed within the community). The second measure, the ‘assortativity index’, quantifies the correlation between the node degree of microbes with the node degree of metabolites.

The total number of vertices *v*_*j*_ for *n* microbes *B*_*i*_, and *m* metabolites *M*_*k*_, is *l* = *n* + *m* (with *j* = *i* + *k*). The normalized entropy is calculated using the standard Shannon index^37^. The normalized entropy is given by: *E*_*R*_ = (− Σ_*i*=1_ *v*_*i*_ log *v*_*i*_) /*E*_*M*_, with *E*_*M*_ = − Σ_*i*_(*i*/ Σ_*i*_ *v*_*i*_) log(*i*/ Σ_*i*_ *v*_*i*_). Note that in this case, the index *i* refers to the number of vertices representing the microbial populations (not the metabolites). If there are *μ* auxotrophy-causing mutations in the community, then there will be *s* = *nxm* − *μ* links in the network.

Mathematically, the assortativity is given by a correlation coefficient^38^ defined by:

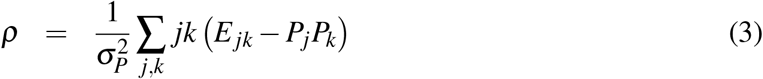

which runs from *−*1 for completely disassortative behavior to 1 for completely assortative. Here, *P*_*k*_ is the normalized distribution of the remaining degree - the number of edges leaving the node, other than the one that connects the pair -, 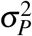 is its variance, and *E*_*jk*_ is the joint probability distribution of the remaining degrees of the two vertices at either end of a link. Note that the network describing auxotrophies links microbes with metabolites and is bipartite, i.e., there are links connecting only microbes to metabolites and *vice versa*, but not microbes to microbes or metabolites to metabolites. Therefore, the relevant description of the interaction is given by a *biadjacency matrix ψ*_*i,k*_.

### 3.3 Cooperative cross-feeding networks

Loss-of-function mutations can cause auxotrophies and thus affect the production of shared metabolites. In addition, other mutations can result in an increased production of shared metabolites within a microbial community. This can occur, for example, by mutations that redirect fluxes within metabolic networks or deregulate biosynthetic pathways^39,18^. As a consequence, the microbial community is comprised of a mixture of auxotrophs and cooperative cross-feeders for different metabolites. We define a parameter *ξ*, denoting the degree of cooperative cross-feeding in the community. This parameter can take values in the range 0 and 1, with 1 indicating that all (100%) metabolites in the given community are produced in increased amounts.

## 4 Results

### 4.1 Model of cross-feeding networks

Our main goal is to understand how environmental disturbance affects different networks of microorganisms that exchange essential metabolites with each other. Specifically, we aim at identifying the parameters that confer robustness against this disturbance. To achieve this goal, we devised a population dynamics model, which describes both the dynamics of all microbial strains that are part of the interaction network and the metabolites that are exchanged between them. The resulting interaction networks can be depicted as a bipartite graph including both the interacting microbes as well as the exchanged metabolites (Fig.1a). Strains within this network can either be able to produce all metabolites they require for growth (i.e., prototrophic genotypes) or lack the ability to produce one or more metabolites (i.e., auxotrophic genotypes). By distributing a certain number of auxotrophy-causing mutations among microbial strains, interaction networks were randomly generated that differed in their size (i.e., number of interacting genotypes) and topology (i.e., distribution of metabolic fluxes among cells). In addition, the amount of metabolites a given microbial genotype produces can be increased to assess how the presence of cooperative phenotypes affects the robustness of the network interaction to ecological disturbance. The resulting microbial networks were exposed to increased levels of disturbance until the population went extinct (Fig.1b,c). The lowest disturbance value, at which a population collapsed, was used to dene the robustness of the focal microbial system.

### 4.2 Increasing numbers of metabolic auxotrophies decreases network robustness to ecological disturbance

To verify how different degrees of metabolic auxotrophies affect the ecological stability of a network of cross-feeding microbes, we randomly introduced auxotrophy-causing mutations at the community level and evaluated the robustness of the resulting microbial system to ecological disturbance.

Our analysis revealed that communities with a higher number of auxotrophy-causing mutations were - on average - less robust to ecological disturbance than communities with a lower number of auxotrophies (Fig.2). The relationship between robustness and the number of auxotrophy-causing mutations in the community can be described with an exponential decay model. Such a model describes the average robustness for each number of auxotrophy-causing mutations (Fig.2). One source of variance at this level is caused by the fact that not all topologies resulting from the randomization process gave rise to microbial systems with a stable equilibrium (i.e., where all microbes and all metabolites are present). Moreover, the variance increased for larger microbial systems (compare Fig.2 a with Fig.2 b), because the number of combinations, in which the auxotrophy-causing mutations can be distributed within the networks (i.e., the number of network topologies), is larger.

**Fig. 2.**
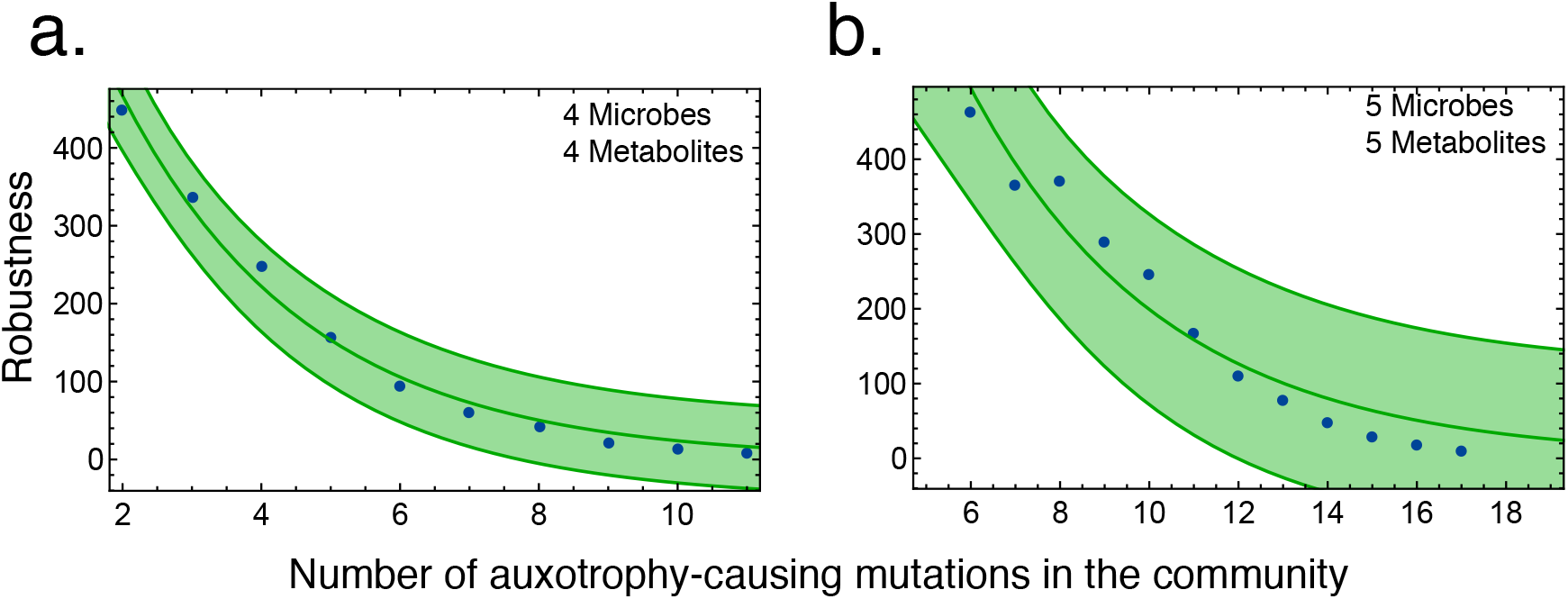
Increasing numbers of auxotrophy-causing mutations impair the robustness of microbial interaction networks to environmental disturbance. The average robustness of interaction networks consisting of **(a)** four microorganisms producing four metabolites, or **(b)** five microorganisms producing five metabolites is shown. Blue points indicate the average robustness for each number of auxotrophy-causing mutations. An exponential decay model is fitted to the data (green line). The 95% confidence interval is shown with a green area.

Together, these results show that auxotrophy-causing mutations can be detrimental for microbial communities, by making them more vulnerable to ecological disturbances.

### 4.3 The topology of auxotrophic networks affects their robustness to ecological disturbance

In our model, we generated different auxotrophic networks by randomly introducing loss-of-function mutations into a given microbial system, which affected the ability of the corresponding microorganisms to produce certain metabolites As a consequence of this procedure, a given microorganism can carry more than one auxotrophy-causing mutation, thus being unable to produce several metabolites simultaneously. Above, we studied the effect of auxotrophy-causing mutations on the robustness of the entire microbial community to ecological disturbance. However, for a fixed number of auxotrophy-causing mutations, several patterns can emerge, depending on how these mutations are distributed among microbes. The range of patterns that can result when mutations are differentially assigned to microorganisms includes cases with a homogeneous distribution of auxotrophy-causing mutations as well as heterogeneous distributions, where some auxotrophs bear the majority of mutations, while all other cells only carry a few.

Our analysis identified two measures, which capture essential properties of the focal topology and correlate with the network robustness to ecological disturbance: (1) *normalized entropy* and (2) *assortativity* (see the “Methods” section for more details) (Fig.3). Simulations show that both normalized entropy and assortativity were positively correlated with robustness to ecological disturbance (Fig.3 c, d). This means that microbial networks, in which metabolite production is more homogeneously distributed, are more robust to ecological disturbance. This is due to the effect the network topology has on the distribution of the microbial population sizes at equilibrium. Asymmetries in the number of metabolites produced by auxotrophs generate an increase in the variance of the distribution of the microbial population sizes at equilibrium, with some populations being present at a lower population frequency, thus making them more prone to extinction. The extinction of one microbe can then trigger a cascade of extinctions of other members in the consortium.

**Fig. 3.**
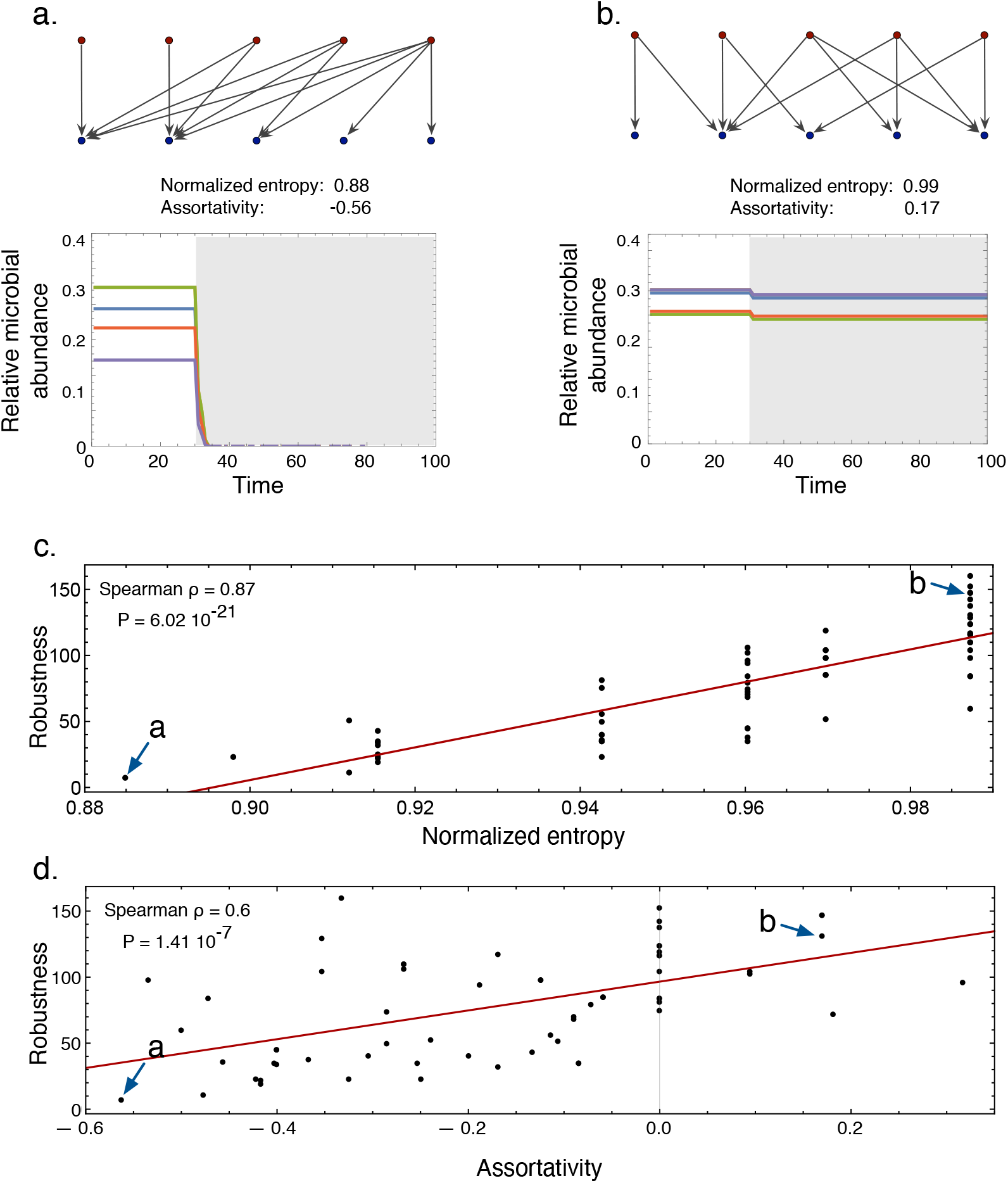
The topology of an interaction network determines its robustness to ecological disturbance. **(a, b)** Two networks consisting of 5 microorganisms with the same number of auxotrophy-causing mutations (here: 12) that exchange 5 different metabolites, yet differ in their network topology, are differentially robust to the same degree of ecological disturbance (*D* = 7). **(a)** The community goes extinct. **(b)** The population is maintained at an alternative stable state. **(c, d)** Statistical relationship between network robustness and **(c)** normalized entropy or **(d)** assortativity. Data represents a microbial system of 5 microbes with 12 auxotrophy-causing mutations in the community that exchange 5 metabolites in total. Arrows point to the networks shown in Fig.3 **(a, b)**.

Thus, our results show that the way auxotrophy-causing mutations are distributed among members of a cross-feeding community (i.e., its topology) strongly affects the robustness of the corresponding communities to ecological disturbances.

### 4.4 Metabolite overproduction increases network robustness to ecological disturbance

So far, we have assumed that mutations in microorganisms only affect their ability to produce certain metabolites. However, mutations may also increase the amount of metabolites a given cell produces^18^. If the resulting metabolite overproduction is costly to the producing cell and the resulting mutant is stabilized by natural selection, the mutation would have transformed the interaction, which was previously based on an exchange of metabolic by-products, into a truly cooperative interaction. However, it is not clear how the presence of such cooperative cross-feeding mutants within a network of auxotrophic cells affects the robustness of the network to ecological disturbance. To test this, we created networks with a different number of auxotrophy-causing mutations and then allowed some fraction of cells to increase the production levels of the remaining metabolic capabilities by a certain magnitude. In all cases, microbes carrying a mutation causing metabolite overproduction payed a fitness cost, which reflected the increased metabolic and energetic investment.

Our results show that the combination of both types of mutations (i.e., auxotrophy-causing and overproduction-causing mutations) can generate networks with a variable response to ecological disturbance. Specifically, the presence of mutations causing metabolite overproduction in an auxotrophic network can increase the robustness of the entire community to ecological disturbance (Fig.4). Networks that were more robust to ecological disturbance emerged when the auxotrophy-causing mutations generate topologies where metabolite production is homogeneous, while the relative position of the mutations causing metabolite overproduction can result in an increase in the abundance of metabolites that are produced by a low number of microbes. Strikingly, some particular combinations of mutations were more robust to ecological disturbance than an entirely prototrophic network (Fig.4 b).

**Fig. 4.**
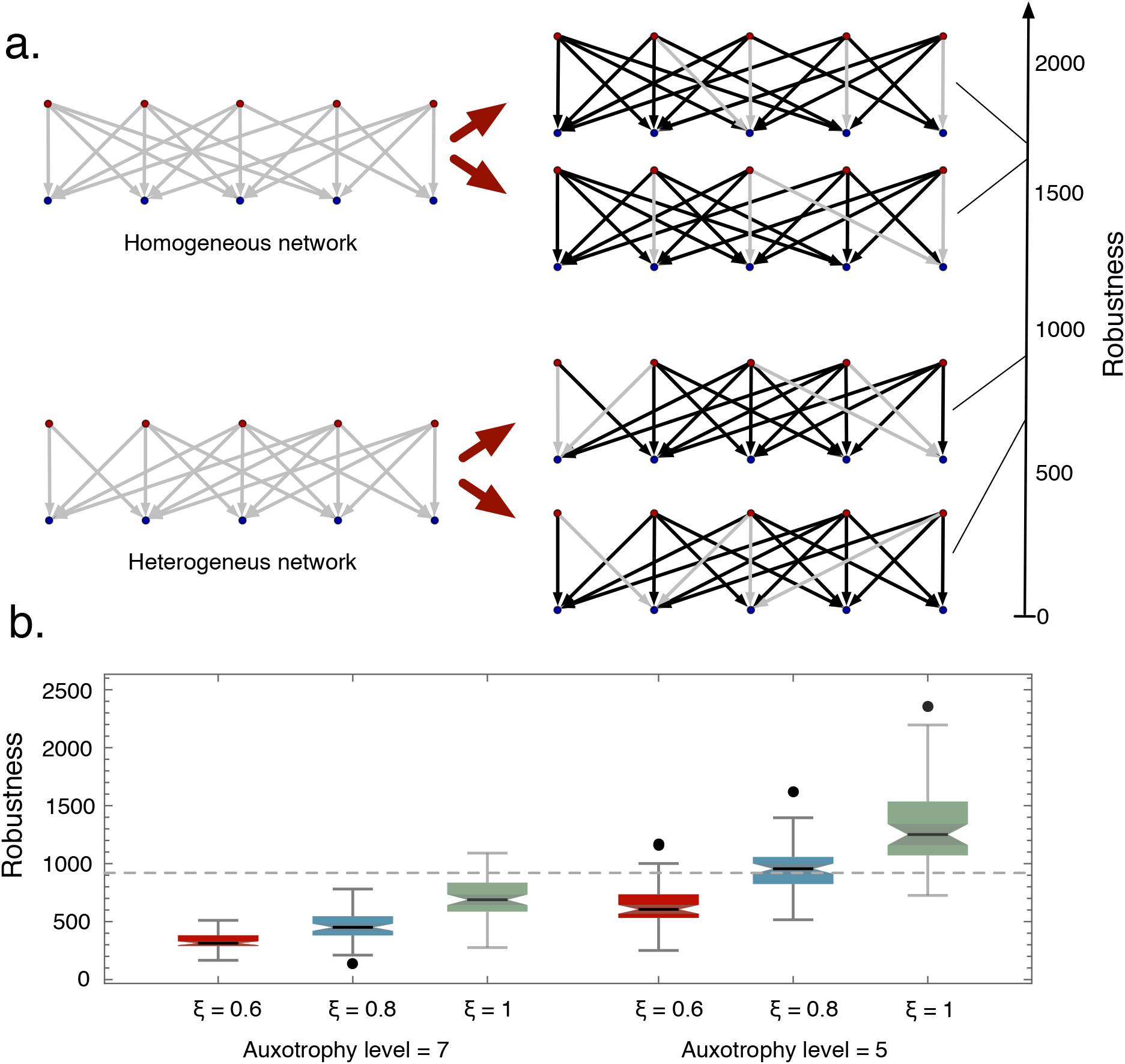
Increased production levels of the public good enhance the stability of interaction networks to environmental disturbance. **(a)** Homogenous and heterogeneous networks can differ in their robustness, depending on which metabolites are produced in increased amounts. In this example, all networks include 5 auxotrophy-causing mutations and 16 links that are being overexpressed (i.e., the degree of cooperative cross-feeding in the community *ξ* = 0.8, black arrows in the network). Increasing the number of links that represent metabolites and which are produced in large amounts, enhances the robustness of the corresponding ecological networks to environmental disturbance. **(b)** Summary of networks with different topologies and different positions of overexpressed links. The dashed line is the robustness level of a prototrophic community (i.e., which does not contain any auxotrophic mutants).

Together, these results show that cooperative cross-feeding networks containing both auxotrophy-causing and overproduction mutations can be highly robust to ecological distrurbance, which may even exceed the stability of prototrophic communities.

## 5 Discussion

In this study, we examined the effect of ecological disturbance on the stability of microbial communities using a dynamical model that describes the interaction between auxotrophic microorganisms and the public goods they produce. This interaction was formalized as a network, which was quantitatively analyzed using different measures of network topology. Our results revealed that (1) communities with more auxotrophy-causing mutations were less robust to disturbance, (2) microbial networks, where the production of public goods was more homogeneously distributed among community members, were more robust to ecological disturbance than networks with a more heterogeneous distribution, and (3) mutations that increased metabolite production levels of auxotrophic microbes within interaction networks increased the robustness of these communities to ecological disturbance.

In the core mathematical model, we assumed that i) disturbance remains constant during the simulation (i.e., press disturbance), ii) auxotrophic microorganisms essentially depend upon an external supply of the required metabolite(−s) to grow (i.e., if one of the essential metabolites cannot be produced anymore by at least one other strain, the whole community collapses), and finally, the metabolites are not affected by the disturbance itself (e.g., metabolites do not degrade). However, relaxing these assumptions does not affect any of our main conclusions

First, if the disturbance stops before the community collapses (i.e., pulse disturbance), the microbial community could still be able to recover and return to the equilibrium. Thus, as a rule of thumb, pulse disturbance is expected to be less disruptive than press disturbance for a given fixed magnitude of disturbance. Moreover, if a set of networks is ordered from more robust to less robust when press disturbance is modeled, the same order is expected to be maintained when pulse disturbance is modeled instead. As a consequence, and given that we are interested in the robustness of a group of networks defined as the magnitude of disturbance leading to the collapse of the whole community, we focused our attention on the case of press disturbance. A more complex, non-trivial behavior is expected to emerge if the microbial system is exposed to sequential pulses of disturbance. However, this issue should be addressed in future studies. Additionally, it is important to mention that a common approach used to determine the stability of a system involves studying departures from the system’s equilibrium when the initial conditions are changed. In our case, this is represented by the initial number of metabolites and the population sizes of the microbial populations in the community. In this way, the stability of microbial communities is modeled as resilience (i.e., the capacity to return to the equilibrium after a transient change in the population size). This approach implicitly assumes that changes in the environment (e.g., pH, temperature, concentration of antibiotics or toxins in the medium) will eventually result in a decline of the population size. In such a framework, cooperation destabilizes microbial networks^22^. However, environmental perturbations of this type are more accurately described by modifying the rate at which microbes replicate (i.e., their Darwinian fitness). Also, the empirical observation of an efficient division of labor between microbes for metabolite production^18,24^ has been ignored in models studying the stability of cooperating microbial networks^22^. By explicitly incorporating perturbations affecting the rate at which microbes replicate in combination with an efficient division of labor for metabolite production into our model, we showed - in stark contrast to a previous study^22^ - a positive effect of metabolic cooperation on the stability of microbial interaction networks. This result emerges, because an enhanced production of the metabolites, which are exchanged among microorganisms, results in a stronger growth response that ultimately increases the robustness of a given community to environmental disturbance.

Second, metabolic interactions among different microbial cells can be obligate or facultative for growth and reproduction of the metabolite-receiving cell. As previously mentioned, we assumed interactions to be obligate, which would for example be the case of auxotrophic bacteria that lack the ability to autonomously produce certain amino acids. However, public goods might not be essential, but can still significantly contribute to microbial fitness. This would be the case for metabolites such as amino acids, vitamins, or nucleotides that are opportunistically consumed as nutrients, whenever they become available in the environment. Also, enzymes that break down extracellular proteins (i.e., proteases^40^) or sugars (e.g., invertases,^41^) liberate publicly available compounds that can enhance the fitness of other community members. Other public goods that could be involved in facultative interaction networks are secondary metabolites that are produced to repel competitors^42^, deter predators^43^, or kill and degrade prey organisms^44^. Relaxing the assumption of an obligate interaction by using an additive Monod growth model did not affect any of our main conclusions (see Supplementary Material). Given that our results can be applied to different types of interaction networks (e.g., obligate or facultative), our results are relevant to a diverse range of microbial systems. These include ecological communities such as intestinal microbiota^45,46^, soil microbiota^47,48^, or microbiota living in aquatic environments^49^.

Third, research shows that modeling mutualistic interactions, without explicitly accounting for the dynamics of resources that mediate interactions between species, can significantly alter conclusions regarding the long-term stability of microbial communities^23^. Given that in our model, we explicitly describe the dynamics of metabolite production and consumption, conclusions regarding the stability will not be altered by these model simplifications present in other studies. In the core model, we have assumed that disturbance only affects the microbial community, yet not the traded metabolites. This would be the case, for example, when the disturbance is due to the presence of an antibiotic in the environment. However, disturbances such as changes in the pH or temperature might affect both the microbial community and the corresponding metabolites themselves (e.g., by changing their chemical properties that affects the chemical’s bioavailability or by chemically degrading the nutrient). Nevertheless, a relaxation of this assumption also yields results that are consistent with the main findings of our study (see Supplementary Material).

The space of potential network configurations is expected to be strongly affected by how accessible the shared metabolites are to other community members. Different factors will have an impact on this, such as a limited diffusion in spatially structured environments^21^, contact-dependent transfer of metabolites via specialized structures^50,51,52^, or a decreased metabolite production by changes in the intracellular metabolic network architecture^53^. Future work should evaluate the role of spatial structure and specialized transport mechanisms for determining the stability of intercellular metabolic networks. Also, the incorporation of metabolic parameters that can be obtained, for example through flux balance analysis^54^, could reveal interesting insights into the dynamics of metabolite exchange within a given microbial community facing disturbance.

Both theory and experiments suggest that microbial population dynamics and the evolutionary dynamics of genes, which are associated with cooperative phenotypes such as the production of public goods, operate on similar timescales and can be linked to each other via an eco-evolutionary feedback loop^55^. As a consequence, a microbial network of auxotrophic mutants may respond to an ecological disturbance by adapting evolutionarily to the corresponding selection pressure. One possible response could be to increase the amount of metabolites that are being overproduced. Indeed, it has been shown previously that metabolite production can change as a phenotypic response to environmental stress^56^. If a microorganism within a given network increased the production of a certain metabolite as a response to the disturbance, this would imply a change in one link of the network. The resulting new network would be more or less robust to the environmental disturbances. In our model, we did not consider a dynamic change in the network as a response to environmental disturbance and instead assumed fixed levels of metabolite production. However, we explored a statistically meaningful and representative subset of the relevant categories, including networks with different degrees of auxotrophy (Fig.2 and Fig.3), different levels of metabolite overproduction, and their combinations (Fig.4). Future work should address how eco-evolutionary feedback loops within microbial auxotrophic networks respond to ecological disturbance.

Here, we have analyzed the ecological stability of intercellular metabolic networks consisting of auxotrophic and cooperative cross-feeding microorganisms. We have identified the amount of public goods that is produced by a given microbial community as well as the way these metabolites are exchanged among community members (i.e., the network topology) as key parameters determining the stability of the whole system. However, the production of metabolites that are being exchanged may not be independent of each other, but can potentially be interconnected through the underlying biosynthetic pathways. Thus, epistatic interactions among mutations causing auxotrophy^57^ and/ or metabolite overproduction may strongly affect how natural selection operates on a given microbial network. Future work should dissect how the topology of intracellular metabolic networks affects the topology of networks that can emerge between cells. Our approach is general, and can therefore be interpreted as a null model for microbial community interactions via public goods in the absence of these constraints. Previous studies analyzed the emergence of cooperative phenotype mainly through the lens of social evolution. By examining the ecological dynamics of different cross-feeding networks, we discovered that increased production levels of exchanged metabolites can significantly enhance the stability of the whole microbial community. Our findings thus provide an intuitive explanation for the evolution and maintenance of metabolic cooperation, suggesting that these cooperative genotypes may be more widespread than previously thought.

## Authors contributions

CK and LO conceived the initial research direction; LO refined concepts, conceived the approach, created and analyzed all models; LO wrote the initial version of the manuscript, which was amended by CK.

## ACKNOWLEDGMENTS

The authors would like to thanks the University of Osnabrück for funding and the Kostlab for fruitful discussions and helpful comments.

