## Supplementary Material for "Cooperation increases robustness to ecological disturbance in microbial cross-feeding networks"

In the following we incorporate several variations to the main model and demonstrate the generality of the results shown in the main text.

### 1 Main model details

$$\frac{dB_i}{dt} = B_i\left(\Gamma(M_k) - \kappa_i B_i - \sum_{\substack{i=1\\ i\neq i}}^{n} \phi_i B_i - D\right) \tag{1}$$

$$\frac{dM_k}{dt} = \sum_{i=1}^{n} \psi_{i,k}\Omega_i B_i - M_k\left(q_k + \sum_{i=1}^{n} d_i B_i - \alpha D\right) \tag{2}$$

Where $\Gamma(M_k) = \prod_{k=1}^{m} M_k\left(r_k - \psi_{i,k}c_k\right)$ in the main model, assuming an obligatory dependence on metabolites, and for a scenario where there is a facultative dependence on metabolites we have $\Gamma(M_k) = \sum_{k=1}^{m}\left(r_k - \psi_{i,k}c_k\right)\mu\frac{M_k}{M_k+K_S}$.

In the first model, the per capita growth of bacteria is the result of the utilisation of all metabolites with $r_k$ denoting the metabolite dependent per capita growth rate, and $c_k$ is the cost associated with metabolite production. The obligatory nature that bacteria have with metabolites is represented by the term $\prod_{k=1}^{m} M_k\left(r_k - \psi_{i,k}c_k\right)$. This product ensures that when at least one of the metabolites $M_k$ is zero, all bacteria will go extinct.

### 2 Extensions of the main model

#### 2.1 Metabolites affected by disturbance

In the core model, we have assumed that disturbance affects only the microbial community, yet not the traded metabolites. However, in several scenarios, disturbances such as changes in the pH or temperature might affect both the microbial community and the corresponding metabolites. To explore the consequences of this, we explored a model where metabolites are also affected by disturbance. An increment in the degree at which metabolites are affected by disturbance ($\alpha$) decrease the robustness to disturbance of the microbial system (Fig.S1).

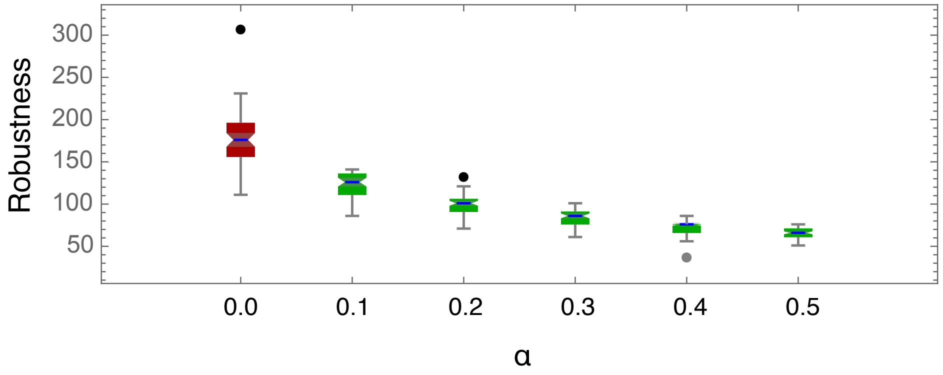

**Fig. S1.** Comparison between the core model (red) with the MA model (Metabolites Affected by disturbance) (green). Here we assumed mutations causing auxotrophy equal to 5, and no mutation causing over-expression (no cooperative cross-feeding), in the plot $\alpha$ value from 0.0 to 0.5. Data represents microbial systems of 5 microbes with 12 auxotrophy-causing mutations in a community that exchanges 5 metabolites in total.

#### 2.2 Additive Monod-type dependence in metabolites: facultative association

In the core model, we have assumed that all metabolites are essential for microbial survival and growth (i.e., interactions are obligate). However, even though several metabolites are not essential, they can still signicantly contribute to microbial tness. Examples of this type of facultative interactions such as the release of vitamins that are opportunistically consumed by other cells as nutrients, extracellular enzymes, which break down extracellular proteins or sugars. To explore if our results are also valid when the interaction between the microbial populations and the metabolites they produce is facultative, we implement this interaction using an additive Monod growth model.

#### 2.3 Testing the main observations for all the model variants

The combination of the extensions introduced in sections 2.1 and 2.2 for the core model generates three different scenarios that we explored in detail in Fig.S2, Table S1 and Fig.S3. The extension where we introduce the possibility of an effect of disturbance on metabolites, is called *MA model*. We called the extension, where we assume a facultative type of dependence of microbes on metabolites "*Monod model*". We found that auxotrophy has a negative impact, reducing robustness in all cases (Fig.S2). We show that correlations between robustness and both normalized entropy and assortativity are significant in all cases (Table S1). Finally, results show that, as it was obtained with the core model (Fig.4, main text), metabolite overproduction increases network robustness to ecological disturbance (Fig.S3).

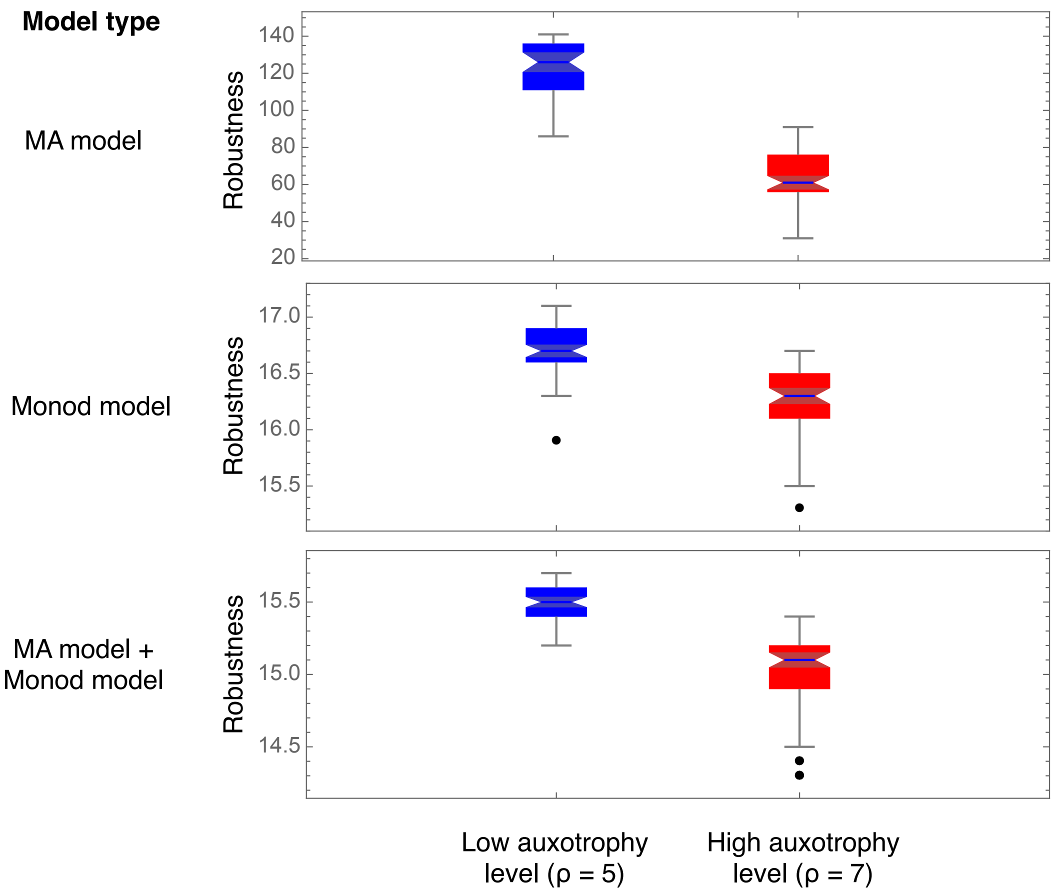

**Fig. S2.** Networks with high a high degree of auxotrophy are less robust to ecological disturbance. Blue color indicates a low auxotrophy level ($\rho = 5$), while red color indicates a high auxotrophy level ($\rho = 7$). Data represents microbial systems of 5 microbes with 5 and 7 auxotrophy-causing mutations in a community that exchanges 5 metabolites in total.

**Table S1:** Network topology affects robustness to ecological disturbance under different model assumptions.

| **Model type** | *normalized entropy* | *assortativity* |
|---|---|---|
| **MA model** | $\rho = 0.62$<br>$P = 1.8\ 10^{-8}$ | $\rho = 0.88$<br>$P = 1.0\ 10^{-22}$ |
| **Monod model** | $\rho = 0.64$<br>$P = 1.7\ 10^{-9}$ | $\rho = 0.9$<br>$P = 8.4\ 10^{-26}$ |
| **MA**<br>**+ Monod model** | $\rho = 0.57$<br>$P = 2.6\ 10^{-7}$ | $\rho = 0.89$<br>$P = 3.1\ 10^{-26}$ |

As shown in the main text (Fig.3), a significant association exists between robustness and different measurements of the homogeneity of networks (measured with the Normalized entropy and Assortativity), for the different model extensions. Data represents microbial systems of 5 microbes with 12 auxotrophy-causing mutations in a community that exchanges 5 metabolites in total.

#### 2.4 Testing the robustness of the model: random parametrization

For all simulations in the main text, we created microbial systems assuming that all parameters affecting the dynamics are the same among microbes and metabolites. Only the position of auxotrophy-causing mutations in the network was varied. This allowed us to study the stability of the resulting network topology of metabolite production in isolation, without the confounding effect that would result if parameters were different among bacteria and metabolites. To ensure that our results still hold when this assumption is relaxed, we assigned random parameters for microbes and metabolites in another set of simulations. Results from randomizing parameters within the ranges shown in Tables S2 and S3, are in line with all the conclusions from the results shown in the main text.

First, 50 random systems with randomized parameters were generated

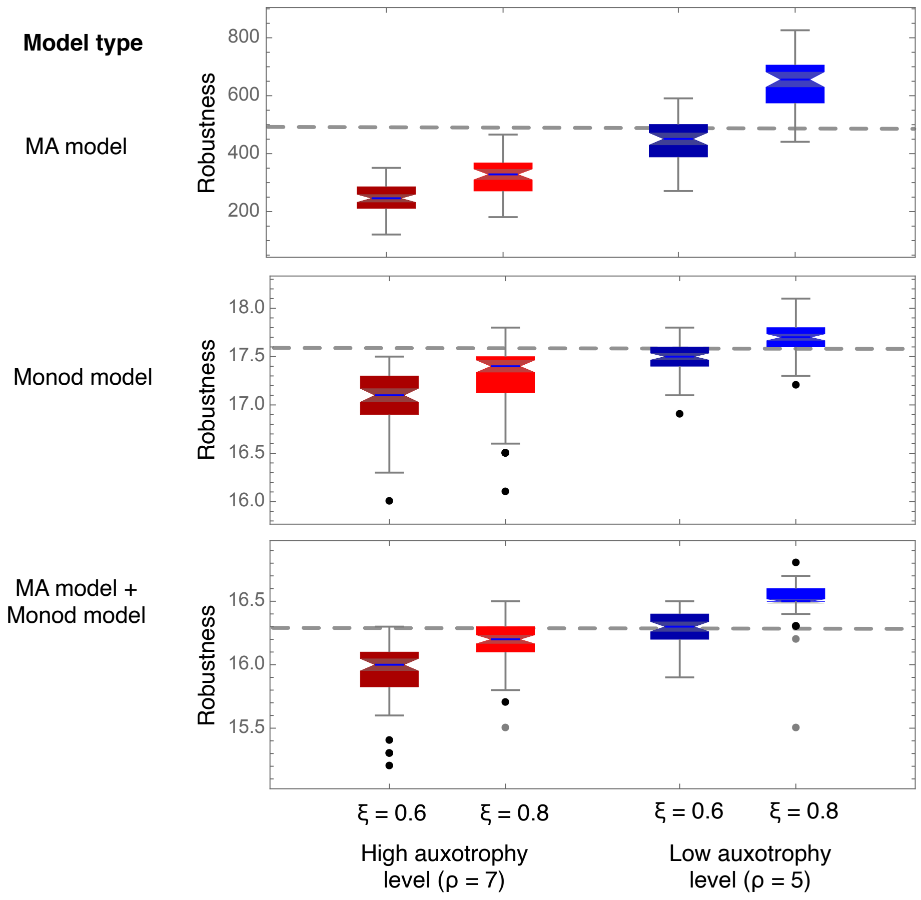

**Fig. S3.** An increase in the production levels of the public goods increases robustness to environmental disturbance in all types of models. Blue color indicates low auxotrophy level ($\rho = 5$), while red color indicates high auxotrophy level ($\rho = 7$). Data represents microbial systems of 5 microbes with 5 and 7 auxotrophy-causing mutations in a community that exchanges 5 metabolites in total.

for each model, resulting in a pattern similar as in Fig.2 from the main text. A log transformation of the data for the robustness resulted in a linear relationship between auxotrophy-causing mutations and the logarithm of the robustness to environmental disturbance. In all cases, there was a negative slope for the regression and a large coefficient of determination ($R^2$) (Fig.S4 a.).

Second, to extend the results obtained in Fig.3 from the main text to systems with randomized parameters, we generated microbial systems with 12 auxotrophy-causing mutations with parameters randomly chosen from the range shown in Tables S2 and S3 and studied the effect of the topology of an interaction network on the robustness to ecological disturbance. In all cases, the *normalized entropy* (blue points), and the *assortativity* (red points), are strongly correlated with robustness (Fig.S4 b.).

Finally, to extend the results obtained in Fig.3 from the main text to systems with randomized parameters, we generated microbial systems with 5 microbes exchanging 5 metabolites, different numbers of auxotrophies (from 5 to 8), and different degrees of cooperative cross-feeding ($\xi$). From all the microbial systems randomly generated, we obtained sets where at least the robustness of one network within a set or, the average robustness of networks within a set was larger than that of a prototrophic community (Fig.S4 c.).

#### 2.5 Effect of the population size on the robustness to ecological disturbance

Asymmetries in the number of metabolites produced by auxotrophs generate an increase in the variance of the distribution of the microbial population sizes at equilibrium, with some populations being present at a lower population frequency, thus making them more prone to extinction. The extinction of one microbe can trigger a cascade of extinctions of other members in the consortium. For the different models we observed a clear link between the robustness of a microbial system and the microbial population with the smallest population size within the community (Fig.S5).

**Table S2:** List of parameters, description, and numerical values explored for the core model.

| **Parameter** | **Description** | **Default value** | **Range** |
|---|---|---|---|
| $r_k$ | Per capita growth rate | 0.9 | 0.7 - 0.95 |
| $\kappa_i$ | Rate of intra-specific competition | 0.2 | 0.05 - 0.35 |
| $\phi_i$ | Rate of inter-specific competition | 0.01 | 0.005 - 0.025 |
| $d_i$ | Rate of metabolite intake | 0.15 | 0.05 - 0.2 |
| $\Omega_i^*$ | Metabolite production rate | [1.05, 1.2 ] | [1.0 - 1.1, 1.15 - 1.3] |
| $q_k$ | Degradation rate of the metabolites | 0.3 | 0.1 - 0.4 |
| $c_k^*$ | Cost of producing a metabolite | [0.05, 0.065 ] | [0.03 - 0.055, 0.06 - 0.075] |
| $\alpha$ ‡ | Effect of the disturbance on the metabolites | [0.0, 0.2 ] | [0, 0.1 - 0.3] |
| $D$ | Disturbance | [0 $\rightarrow$ extinction] | - |

(*) The first value in the pair for parameters $\Omega_i$ and $c_k$ denotes the default value for production and cost, while the second value denotes production and cost when metabolites are overproduced. (‡) In the core model, the value is zero, while it is non-zero when assuming that metabolites are affected by the disturbance (MA model).

**Table S3:** List of parameters, description, and numerical values explored for the Monod model.

| Parameter | Description | Default value | Range |
|---|---|---|---|
| $r_k$ | Per capita growth rate | 0.9 | 0.7 - 0.95 |
| $\kappa_i$ | Rate of intra-specific competition | 0.2 | 0.05 - 0.35 |
| $\phi_i$ | Rate of inter-specific competition | 0.001 | (5 - 25) $10^{-4}$ |
| $d_i$ | Rate of metabolite intake | 0.15 | 0.05 - 0.2 |
| $\Omega_i^*$ | Metabolite production rate | [1.05, 20.05 ] | [1.0 - 1.1, 15 - 25] |
| $q_k$ | Degradation rate of the metabolites | 0.3 | 0.1 - 0.4 |
| $c_k^*$ | Cost of producing a metabolite | [0.05, 0.065 ] | [0.03 - 0.055, 0.06 - 0.075] |
| $\alpha^{\ddagger}$ | Effect of the disturbance on the metabolites | [0.0, 0.2 ] | [0, 0.1 - 0.3] |
| $D$ | Disturbance | [0 $\rightarrow$ extinction] | - |
| $K_S$ | "Half velocity constant" (a Monod coefficient) | 20 | 17 - 23 |
| $\mu$ | Maximum specific growth rate (a Monod coefficient) | 5 | 3 - 7 |

The first value in the pair for parameters $\Omega_i$ and $c_k$ denotes the default value for production and cost, while the second value denotes production and cost when metabolites are overproduced. (‡) In the Monod model, the value is zero, while it is non-zero when assuming that metabolites are affected by the disturbance (Monod + MA model).

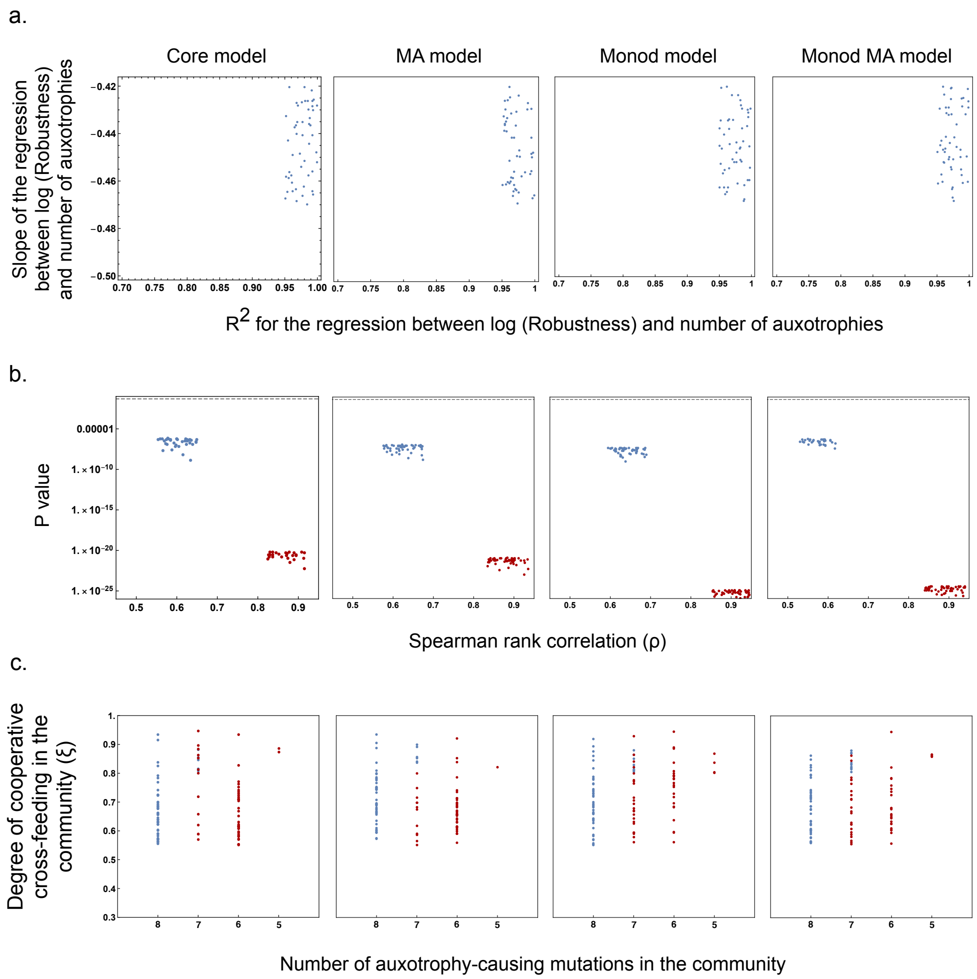

**Fig. S4.** Models with randomized parameters for 5 microbes exchanging 5 metabolites. **a.** Systems resulted in a pattern similar to Fig.2 from the main text. A linear relationship between auxotrophy-causing mutations and the logarithm of the robustness resulted in these systems. In all cases there was a negative slope for the regression and a large coefficient of determination ($R^2$). **b.** Points represent different network topologies of microbial systems with 12 auxotrophy-causing mutations. In all cases the normalized entropy (blue points) and the assortativity (red points), are strongly correlated with robustness. **c.** Lowest number of a) auxotrophy-causing mutations and b) degree of cooperative cross-feeding where, within a set of networks, in at least one network (blue) or on average (red), the robustness was larger than that of a prototrophic community.

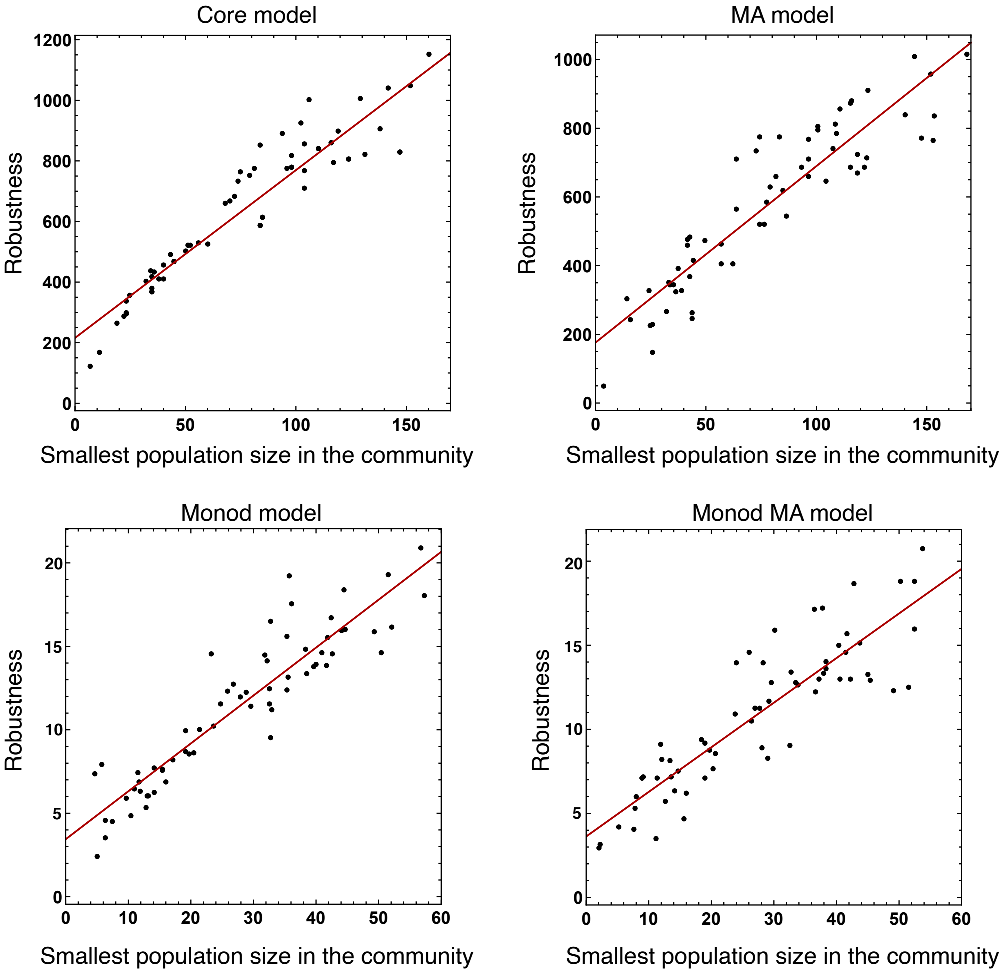

**Fig. S5.** Correlation between robustness and the smallest microbial population size in the community. Data represents microbial systems of 5 microbes with 5 and 7 auxotrophy-causing mutations in a community that exchanges 5 metabolites in total. Parameters where randomized from ranges shown in Tables S2 and S3.
